# Common datastream permutations of animal social network data are not appropriate for hypothesis testing using regression models

**DOI:** 10.1101/2020.04.29.068056

**Authors:** Michael N. Weiss, Daniel W. Franks, Lauren J. N. Brent, Samuel Ellis, Matthew J. Silk, Darren P. Croft

## Abstract

1. Social network methods have become a key tool for describing, modelling, and testing hypotheses about the social structures of animals. However, due to the non-independence of network data and the presence of confounds, specialized statistical techniques are often needed to test hypotheses in these networks. Datastream permutations, originally developed to test the null hypothesis of random social structure, have become a popular tool for testing a wide array of null hypotheses. In particular, they have been used to test whether exogenous factors are related to network structure by interfacing these permutations with regression models.
2. Here, we show that these datastream permutations typically do not represent the null hypothesis of interest to researchers interfacing animal social network analysis with regression modelling, and use simulations to demonstrate the potential pitfalls of using this methodology.
3. Our simulations show that utilizing common datastream permutations to test the coefficients of regression models can lead to extremely high type I (false-positive) error rates (> 30%) in the presence of non-random social structure. The magnitude of this problem is primarily dependent on the degree of non-randomness within the social structure and the intensity of sampling
4. We strongly recommend against utilizing datastream permutations to test regression models in animal social networks. We suggest that a potential solution may be found in regarding the problems of non-independence of network data and unreliability of observations as separate problems with distinct solutions.

## Introduction

Social structure, defined as the patterning of repeated interactions between individuals (Hinde 1976), represents a fundamental characteristic of many animal populations with far-reaching consequences for ecology and evolution, including for gene-flow, social evolution, pathogen transmission, and the emergence of culture (Kurvers et al., 2014). The last two decades have seen widespread adoption of social network methods in animal behaviour research to quantify social structure (Webber & vander Wal, 2019). The network framework is appealing because it explicitly represents the relationships between social entities from which social structure emerges (Hinde, 1976), and thus allows tests of hypotheses about social structure at a variety of scales (individual, dyadic, group, population). Social networks can be based on direct observations of interactions, or inferred from other data types, such as groupings of identified individuals (Franks et al., 2010), GPS tracks (Spiegel et al., 2016), proximity loggers (Ryder et al., 2012), or time-series of detections (Psorakis et al., 2012).

The analysis of animal social network data presents a statistical challenge. Specifically, two separate issues must be addressed. First, network data are inherently non-independent, thus violating the assumptions of independent observations inherent to many commonly used statistical tests. Second, factors outside of social structure, such as data structure and observation bias, may influence the structure of observed animal social networks, potentially leading to both type I and type II errors in statistical tests (Croft et al., 2011).

To address the problem of non-independence, a wide array of statistical tools have been developed, primarily in the social sciences. These methods include permutation techniques that allow for hypothesis testing in the presence of non-independence. These permutations normally test relationships between exogenous variables and network properties, such as the presence and strength of social ties, or the centrality of nodes in the network. These methods typically build empirical null distributions by randomly assigning the location of nodes in the network, while holding the network structure constant (“node-label permutations”). The resulting null distribution maintains the non-independence inherent to the network while breaking any relationship that exists between network structure and potential covariates (Dekker et al., 2007).

While these methods are useful for dealing with the issue of non-independence, they do not address the second issue, from which studies of animal social systems in particular often suffer. Because the methods developed in the social sciences only permute the final constructed network, they do not inherently account for common biases in the collection of the raw observational data used to construct the final network. These biases may be introduced by the method of data collection (e.g. group-based observations), individual differences in identifiability, or demographic processes (James et al., 2009). For example, consider a situation where researchers are interested in differences in social position between sexes, but females are more cryptic and thus observed with a lower probability. This would lead to incorrect inferences due to biases in the observed network structure that are unrelated to the true social processes of interest (Farine, 2017). To deal with these problems, a suite of alternative permutation procedures has been developed. Rather than permuting the final network, these methods permute the raw data used to construct the network. These methods are therefore sometimes referred to as “datastream permutations.” The goal is to construct permuted datasets that maintain structures of the original data that may influence the observed network structure (e.g. the number of times individuals were observed and the sizes of observed groups), while removing the social preferences that underpin the social network (Farine & Whitehead, 2015).

The original datastream permutation technique for animal social data was proposed by Bejder et al. (1998), based on the procedure outlined by Manly (1997) for ecological presence-absence data. Bejder et al.’s procedure was designed to test whether a set of observed groupings of identified animals showed signs of non-random social preferences. This procedure permutes a group-by-individual matrix, where rows are groups and columns are individuals, with 1 representing presence and 0 indicating absence. The algorithm finds 2 by 2 “checkerboard” submatrices, with 0s on one diagonal and 1s on the other, that can be “flipped” (i.e. 0s replaced with 1s and vice versa). These flips maintain row and column totals (the group size and observations per individual, respectively), but permute group membership. In biological terms, matrices generated with this procedure represent the null hypothesis that individuals associated completely at random, given the observed distribution of group sizes and the number of sightings per individual.

Refinements of this method were later developed that constrained swaps within time periods, classes of individual, or locations (Whitehead et al., 2005). One alteration also controls for gregariousness, and allows for permutation of association data not constructed using group membership (Whitehead, 1999). Controlling for gregariousness and sighting history is possible when each sampling period is represented as a square matrix, where 1 indicates that individuals associated in that period and 0 indicates no association. In this format, the data can be permuted in a way that maintains the number of associates each individual had in each sampling period (Whitehead, 1999).

In recent years, datastream permutation methods have been developed that can handle more complex data structures, such as GPS tracks (Spiegel et al., 2016), time-series of detections (Psorakis et al., 2015), and focal follow data (Farine, 2017). All of these methods have in common that they essentially randomise raw observations of social association (or interactions) data and thus remove social structure while maintaining most other features of the data, including features potentially causing biased measurements of social structure. They thus provide a robust null distribution to test for non-random social structure in a dataset, which is a key step in understanding the behavioural ecology of wild populations.

Many empirical studies and methodological guides have suggested interfacing these null models with other statistical techniques, particularly regression models (including ordinary least squares, generalized linear models, and mixed-effects models), to test hypotheses about network structure. The logic of this recommendation is that permutation-based null models allow researchers to account for sampling issues when testing hypotheses using these common statistical models. However, it is important to recognize the limitations of this approach, and to think carefully about the null hypothesis that these methods specify. In common datastream permutation null models, the null hypothesis specified is that the population’s social structure is random, once we control for the structure of the data and other confounds. For a particular quantity of interest, such as edge weights, node centralities, or differences between networks, this null hypothesis can be equivalently stated as proposing that all variance in a given value or network metric is due to data structure, confounds, and residual variance. In network terminology, this null hypothesis is a random network, within a set of constraints. This null hypothesis is by design, because this form of permutation procedure was specifically created to test for non-random social structure. However, we feel there has been a lack of consideration about whether this null hypothesis is appropriate in other contexts, such as regression modelling. We show here that these procedures do not provide an appropriate null hypothesis for testing the null hypotheses of regression models.

Consider the basic linear model: 

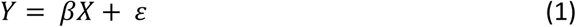

where *Y* is a response variable, *X* is a matrix of predictor variables, *ε* is the error term, and *β* is a vector of estimated coefficients. We are typically interested in testing the null hypothesis *β* = 0, representing no relationship between the response *Y* and the predictor(s) *X*. In permutation based hypothesis testing procedures, researchers specify this null distribution by randomising either *X* or *Y*, often with constraints, thus maintaining the distribution of values but breaking any covariance between the variables (Anderson & Robinson, 2001). This is the logic behind traditional node-label permutation tests of regression in social networks (Croft et al., 2011).

Datastream permutations, however, do something very different, which is inappropriate for testing this hypothesis. By permuting the data underlying network measures and then re-calculating the response variable, these procedures change the distribution of *Y*, instead of breaking relationships between the variables. If the network has non-random social structure, even structure entirely unrelated to *X*, then we will typically see a reduction in the variance of *Y* as we permute the raw data. When *Y* has a larger variance in the observed data than in the permutations, more extreme values of *β* are more likely to occur in the observed data, even if the null hypothesis is true. This procedure is therefore likely to result in much higher rates of false-positive type I error than is acceptable (Figure 1).

**Figure 1.**
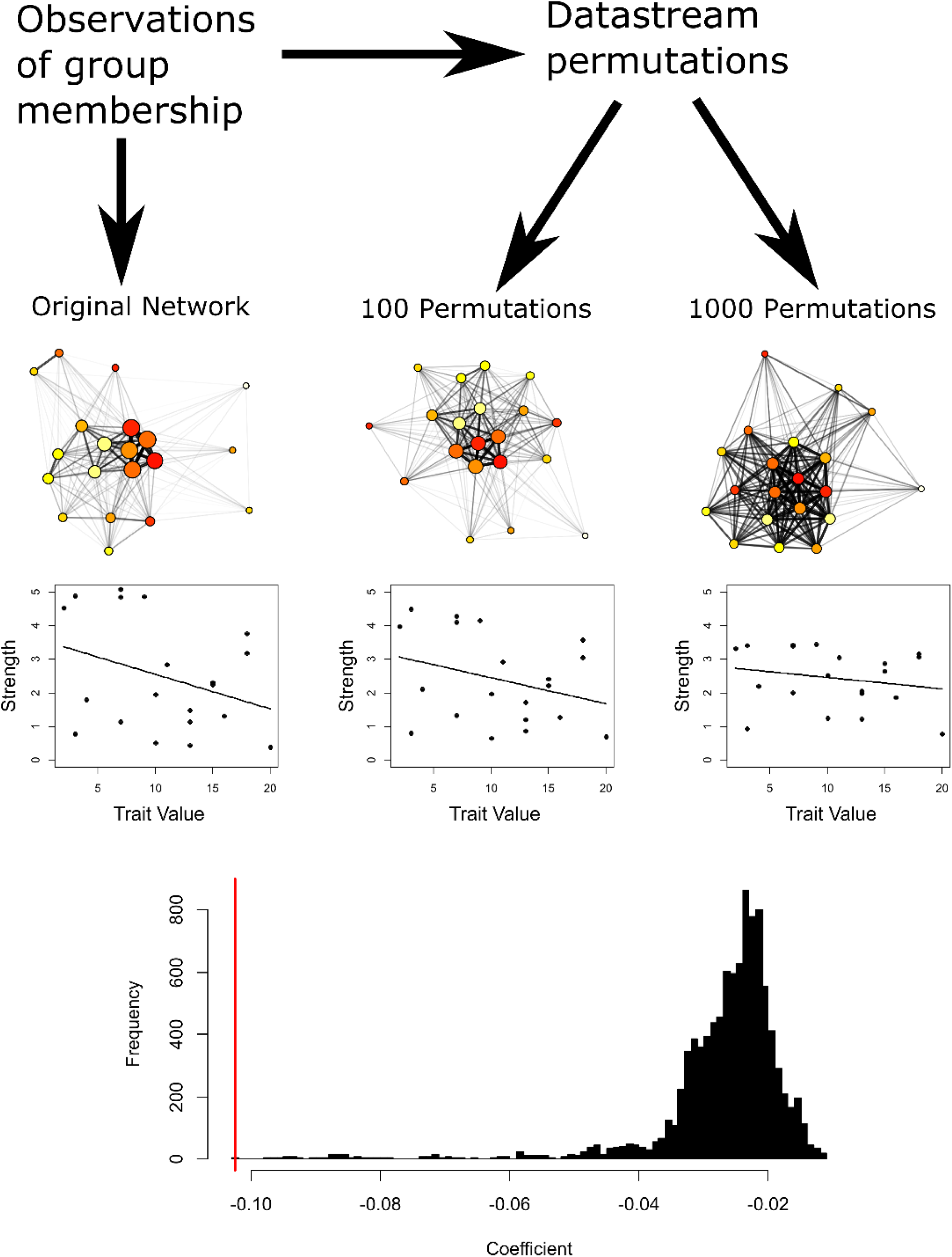
Example of the mechanism by which datastream permutations may lead to false positives in linear regression. In the original network, there is variation in strength among individuals driven by differences in gregariousness (represented by node size in the social networks). Individuals are assigned a trait value (represented by colour in the social network) unrelated to their network position. By chance, there is a slight negative relationship between network strength and trait value in the observed network. After several permutations, there is a reduction in the variance in the strength of individuals in the permuted network, and thus the magnitude of the relationship is reduced. The bottom histogram shows the distribution of null coefficients after 10,000 permutations (black), and the coefficient from the original linear model (red).

Changes in variance between the observed and permuted data is more than just a technical issue. There is a fundamental problem with this approach when it comes to testing hypotheses using regression models. When researchers fit regression models to predict network properties from exogenous variables, the null hypothesis they will be testing against can be stated as “the variation in network structure is not related to the exogenous variable.” This, however, is not the null hypothesis tested by the commonly used datastream permutation methods. Rather, the null hypothesis that is proposed by these datastream permutations could be stated as “the degree of variation in network structure and its relationship to the exogenous variable are both due to random interactions of individuals within constraints.” The researcher cannot disentangle the null hypothesis of no relationship between the network and the predictor from the null hypothesis of random social structure. In other words, a significant result from this procedure could be due to a relationship between the predictor and the network, or because individuals do not interact at random, whether or not the true social structure is related to the predictor. This fundamental mismatch between the null hypothesis of interest and that tested by the datastream permutation algorithm makes tests of regression models using this procedure nearly uninterpretable.

Here, we demonstrate the problems that occur when combining datastream permutations of animal social network data with regression using two simulated scenarios. In these scenarios, we generate datasets with simple non-random social structure. We then introduce a random exogenous variable that has no relationship to social structure, and test for a relationship between network structure and this variable with linear models, using datastream permutations to determine statistical significance. We show that even in the absence of any true relationship between exogenous variables and social structure, datastream permutations are highly prone to producing significant *p*-values when social structure is non-random. We caution against using these datastream permutations to test the coefficients of regression models, and we discuss possible solutions and alternative methods for regression analysis in social networks.

## Methods

### General framework

To illustrate the problems with using datastream permutations to test the coefficients of regression models, we carried out simulations across two different scenarios, reflecting common research questions in animal social network analysis. The first scenario simulates a case in which researchers are interested in whether dyadic covariates (e.g. kinship or phenotypic similarity) influences the strength of social bonds, which we will refer to as a case of “dyadic regression”. The second scenario simulates a case when researchers are interested in how a quantitative individual trait (e.g. age or personality) influences individual network position, which we refer to as “nodal regression.”

While the methods of network generation differ slightly for each scenario, the general steps are the same:

1. Generate observations of a network in which the quantity of interest (edge weight or node centrality) has inherent variation.
2. Generate values for a trait that are unrelated to this variation.
3. Fit a linear model with the network property as the response variable and the trait as the predictor
4. Create permuted versions of the observed network via a common datastream permutation
5. Compare the original model’s coefficient to those fit to the permuted data to calculate a *p*-value

For each simulation, we perform 200 runs, with varying parameter values (Table 1). For each run of both simulations, we produce four outputs. The first two outputs are the coefficient of the fitted linear model and the *p*-value from the permutation test. The other two outputs give information about the characteristics of the dataset. The first is the standard deviation of the response variable (either the edge weights or weighted degrees), indicating the degree of non-randomness in the social structure, and the second is the average number of sightings per individual, a common measure of sampling effort in social network studies.

**Table 1.** Ranges for varied parameters used in simulations

| Parameter | Meaning | Dyadic | Nodal | Range |
| --- | --- | --- | --- | --- |
| $N$ | Number of individuals in population | ✓ | ✓ | 20 – 100 |
| $\mu$ | Mean association probability | ✓ | | 0.01 – 0.5 |
| $t$ | Number of sampling periods | ✓ | | 20 – 200 |
| $\phi$ | Precision of beta distribution for association probabilities | ✓ | | 1 – 10 |
| $o$ | Observation probability per sampling period | ✓ | | 0.1 – 1 |
| $G$ | Number of observed groupings | | ✓ | 20 – 500 |
| $M$ | Maximum grouping size | | ✓ | 5 – 10 |
| $\sigma$ | Standard deviation of group size preference | | ✓ | 0.1 – 2.0 |

### Dyadic regression: Does similarity in a trait predict the strength of social relationships?

In our first simulation, we investigate the case in which the researcher is interested in the influence of a dyadic predictor (such as similarity in phenotype or kinship) on the rates at which dyads associate or interact. Our simulation framework is heavily inspired by those of Whitehead & James (2015) and Farine & Whitehead (2015). We simulate a population of *N* individuals, and assign each dyad an association probability *p*_*ij*_ from a beta distribution with mean *μ* and precision *ϕ* (*α* = *μϕ, β* = (1-*μ*) *ϕ*). By assigning association probabilities in this way, we create non-random social preferences in the network, and thus larger variance in edge weights than would be expected given random association (Whitehead et al., 2005).

We then simulate *t* sampling periods. For simplicity, individuals are sighted in each sampling period with a constant probability *o*, and associations between dyads where both individuals are sighted occur with probability *p*_*ij*_. We then build the observed association network by calculating dyadic simple-ratio indices: 

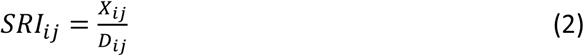

Where *X*_*ij*_ is the total number of sampling periods in which *i* and *j* were observed associating, and *D*_*ij*_ is the total number of periods in which either *i* or *j* was observed (including periods where they were observed, but did not associate with any individuals).

We then assign each individual a trait value from a uniform distribution (0,1). We do not need to specify what this trait represents for our simulation, but it could represent any quantitative trait used as a predictor in social network studies (age, personality, cognitive ability, dominance rank, parasite load, etc.). Note that the trait value is generated after the observations of association and has no influence on any network property.

We then fit the linear model: 

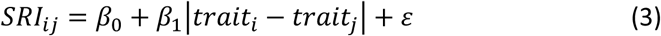

and save the estimate of *β*_1_. We compare this coefficient to a null model generated using the sampling period permutation method proposed by Whitehead (1999). There are several algorithms available to perform these swaps. We use the “trial swap” procedure described by Miklós & Podani (2004) and suggested for social network studies by Krause et al. (2009). For each trial, this procedure chooses an arbitrary 2 by 2 submatrix of the lower triangle within a random sampling period. If a swap is possible, it is performed (and symmetrized), otherwise the matrix stays at its current state. These steps when the matrix is not changed are referred to as “waiting steps.” This algorithm is ideal because it ensures that the Markov chain samples the possible matrices uniformly, while other algorithms that do not include waiting steps exhibit biases in their sampling of the possible matrices (Miklós & Podani, 2004). We generate 10,000 permuted datasets for each simulation, with 1,000 trial swaps between each permutation, and re-fit our linear model to each permuted dataset, recording the coefficient. We then use this distribution of coefficients to calculate the *p*-value of the linear model’s coefficient. Across the 200 runs, we vary the parameters of the simulation by drawing *μ, ϕ, N, o*, and *t* randomly using Latin hypercube sampling (Table 1).

### Nodal regression: Do individual traits influence network centrality?

We next investigate the same concept in the context of nodal regression. This form of analysis tests whether some individual attribute is related to variation in network position. This is perhaps the most common use of datastream permutation null models for testing the significance of linear regression coefficients in animal social networks (e.g. Cowl et al., 2020; Poirier & Festa-Bianchet, 2018; Zeus et al., 2018). For simplicity, we focus on weighted degree, which is simply the sum of an individual’s edge weights.

In this simulation, we consider the case where networks are derived from patterns of shared group membership (“gambit of the group”). This form of data collection is extremely common in animal social network studies, and was the basis for the original datastream null model developed by Bejder et al. (1998).

The framework for this simulation is based on that used by Firth et al. (2017). We simulate *G* observations of groupings in a population of *N* individuals. Each group is assigned a group size *S* from a discrete uniform distribution on [1,*M*]. We assign each individual a preference for a particular group size *P* from a truncated normal distribution with mean (1+*M*)/2, standard deviation *σ*, lower bound 0, and upper bound *M*. Higher values of *σ* will therefore lead to higher variation in gregariousness in the population. For each group *g*, membership is determined by sampling *S*_*g*_ individuals without replacement, with individual sampling probability determined by the size of group *g* and each individual’s group size preference: 

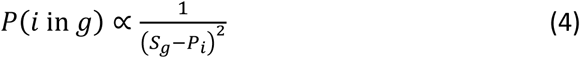

This gives the simulation the property that individuals with higher assigned gregariousness scores tend to be seen in larger groups, and vice versa. This leads to non-random differences in gregariousness (and thus weighted degree) between individuals. We then calculate the association network, again using the SRI: 

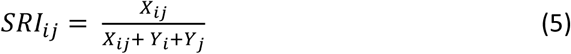

Where *X*_*ij*_ is the number of groups in which the dyad was seen together, and *Y*_*i*_ and *Y*_***j***_ are the number of groups in which only *i* or only *j* were seen, respectively. After calculating the network, we determine each individual’s weighted degree. We again generate a trait value for each individual at random from a uniform distribution on (0,1) and fit the linear model 

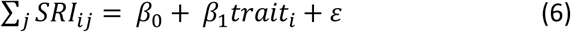

and again save the estimate of *β*_1_. We compare this coefficient to random coefficients fit to networks generated using the group-based permutation procedure proposed by (Bejder et al., 1998). This procedure again sequentially permuted the observed dataset, while maintaining the size of each group and the number of groups per individual. We again use the trial swap method to perform these permutations, generating 10,000 permuted datasets with 1,000 trials per permutation, and derived *p*-values in the same way as above. We vary the parameters of this simulation by using Latin hypercube sampling to draw values of *N, M, G*, and *σ* (see Table 1 for ranges).

### Analysis

We use the outputs of the simulations primarily to derive overall type I error rates for both scenarios, calculated as the portion of runs in which a *p*-value less than 0.05 was obtained. We further investigated the sensitivity of these results to non-random social structure, sampling effort, and population size. Previous work suggests that the sensitivity of datastream permutation techniques are highly dependent on variation in social structure and sampling intensity (Whitehead, 2008). We use binomial generalized linear models to summarize how population size, response variance, and sampling intensity influence the probability of false positives. We further analyse these relationships qualitatively using conditional probability plots.

## Results

### Dyadic regression

The overall type I error rate for the dyadic regression case was high, with 35% of runs giving false positive results (70 out of 200 runs). Sensitivity analysis suggested that the most important factors influencing type I error rate in our simulations were the average number of sightings per individuals and the variance of association probabilities. As the average number of sightings increased, so did the false positive rate (*β* = 0.012 ± 0.004, *z* = 3.149, *p* = 0.002, Figure 2a). Similarly, networks with higher variance in edge weights experienced higher type I error rates (*β* = 8.35 ± 8.93, *z* = 2.37, *p* = 0.02, Figure 2b). There was a less clear, but statistically significant relationship between network size and type I error rates, with networks of larger size typically having lower type I error rates (*β* = - 0.014 ± 0.007, *z* = -2.02, *p* = 0.04, Figure 2c).

**Figure 2.**
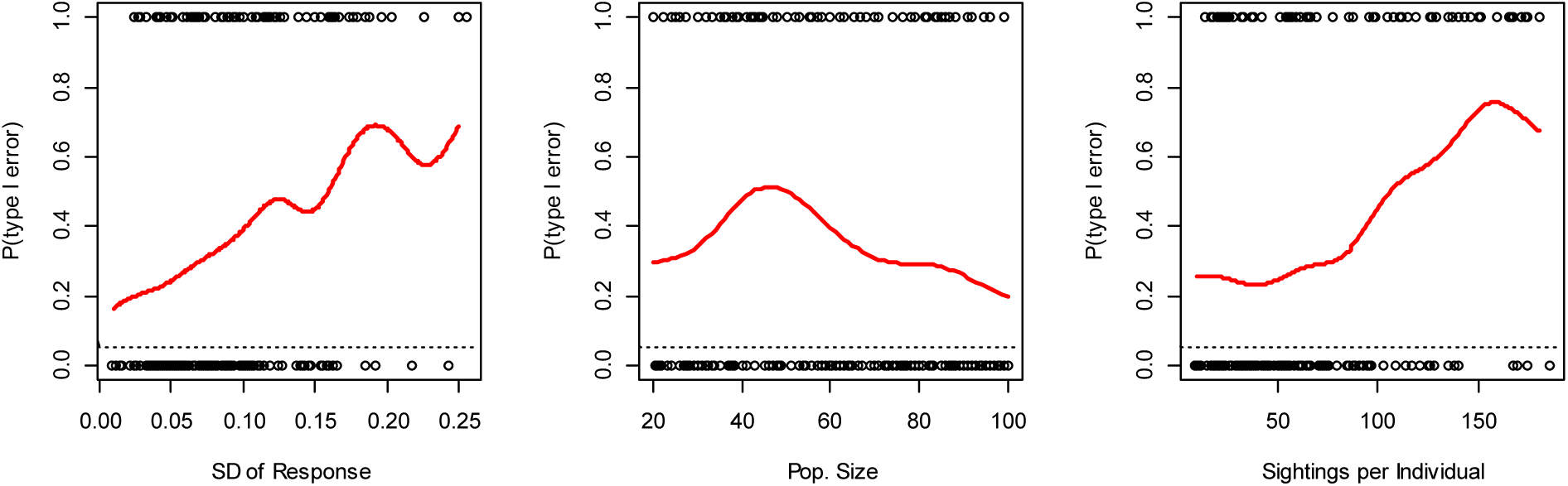
Conditional probability plots from dyadic regression simulation. Points indicate results of individual simulation runs (1 = significant *p*-value, 0 = non-significant *p*-value). Red lines are smoothed condition probabilities of a significant *p*-value. Dotted line indicates target type I error rate of 0.05.

### Nodal regression

The nodal regression case resulted in even higher type I error rates than the dyadic case, with almost half of runs giving false positive results (95 out of 200 runs; 47.5%). The rate of type I errors was strongly influenced by the variance in weighted degree; as the standard deviation of the response increased, so too did the false positive rate (*β* = 1.18 ± 0.50, *z* = 2.34, *p* = 0.019, Figure 3a). In contrast, as the size of the network increased, the false positive rate decreased, although never approaching the target false positive rate of 0.05 in our simulations (*β* = -0.02 ± 0.01, *z* = 2.89, *p* = 0.004, Figure 3c). In this simulation, the number of sightings per individual did not appear to significantly influence the type I error rate (*β* = 0.018 ± 0.013, *z* = 1.43, *p* = 0.153, Figure 3b). This may be because, in networks with few groupings but high sightings per individual, there were fewer possible permutations of the observed network, and therefore the permuted networks were more similar to the original network.

**Figure 3.**
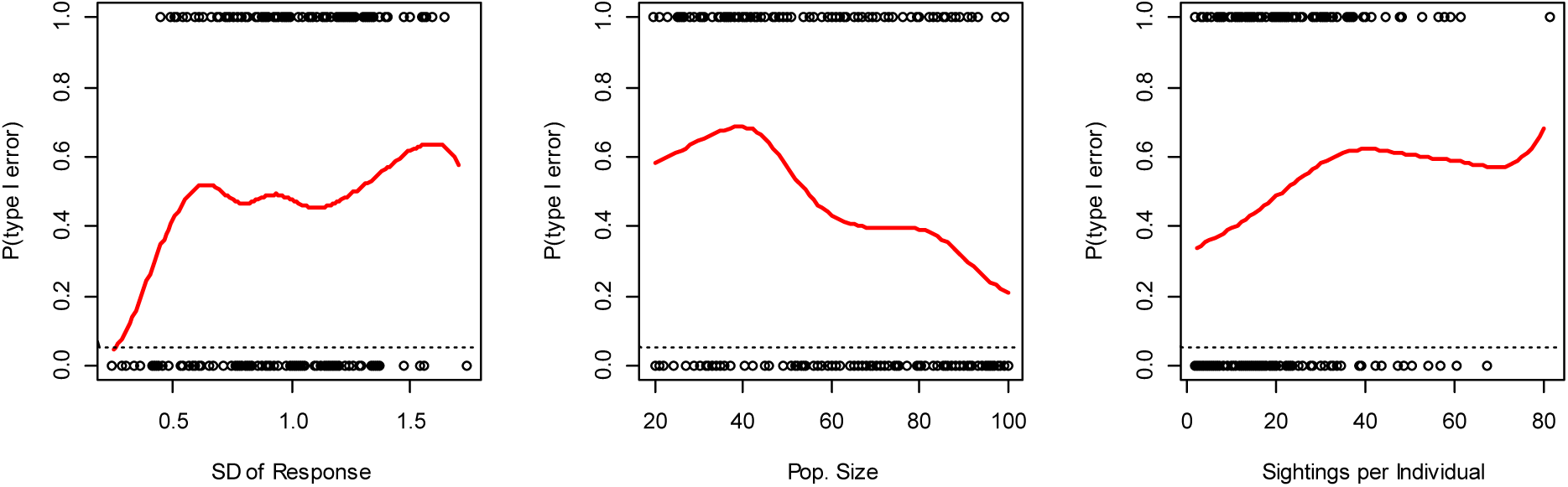
Conditional probability plots of type I error rates for the nodal regression simulation. Points indicate the outcome of individual runs (1 = significant *p*-value at 0.05, 0 = non-significant *p*-value). Red lines are smoothed conditional probabilities of a significant *p*-value. Dotted lines indicate the target error rate of 0.05.

## Discussion

These two simple simulated scenarios show that the commonly used datastream permutation procedures for animal social network data produce extremely high and thus unacceptable false-positive rates when applied to regression models. This is because datastream permutations do not generate appropriate null distributions for testing the significance of model coefficients. We therefore strongly warn against using this procedure.

We now turn to some potential solutions to this problem that may still facilitate inference in these situations. This is not intended to be a comprehensive guide to hypothesis testing in social networks, and other solutions are certainly possible. We encourage other researchers to consider these and other possible solutions.

### Transforming the response variable

If variations in social behaviour are present in the network, datastream permutations undesirably eliminate social influence and reduce the variance in the response of a regression model. A potential fix for this problem is to simply standardize the variable of interest, perhaps to have a mean 0 and standard deviation 1 (*Z*-scores), and to repeat this process for all permutations. While this is likely to reduce type I error rates, we caution that this is a quick fix of the symptom and does not address the cause: that the null model generated does not test the desired hypothesis. We therefore do not consider this to be an adequate solution in itself.

### Alternative test statistics

Another potential solution could be found in using a test statistic other than the coefficients from the model. In the context of node-label permutations such as MRQAP, pivotal statistics such as the *t* statistic for ordinary least squares regression typically perform better than raw coefficient values (Dekker et al., 2007). While previous authors have recommended against using the *t* or *Z* statistic, because they represent deviations from a parametric distribution rather than direct features of the data (Farine, 2017), such statistics could experience a lower type I error rate than those reported here. However, as in the case of transforming the response, this does not address the larger issue of an incorrectly specified null hypothesis. We therefore do not view the adoption of alternative test statistics from regression models compared to datastream permutations as an appropriate solution.

### Separating the issue of non-independence from biases in the data

We suggest that the way forward for hypothesis testing in animal social networks is to recognize that the problems of non-independence of network measures and the influence of data structure underlying networks are separate issues, requiring separate solutions. Not all animal network data will be subject to the issue of unreliability (e.g., in cases where sampling is balanced across subjects and relevant contexts) and in some instances the data may be complete and unbiased. In these cases, node permutations or other statistical network models are appropriate (Croft et al. 2011). In instances where structure in the data needs to be controlled we propose two potential methods; other solutions are certainly possible, and we encourage further work on this matter.

The first method would utilize generalized affiliation indices (GAIs; Whitehead & James, 2015) or similar corrections to account for confounding variables that may influence observed edge weights. GAIs fit the observed data associations or interactions as the response in a binomial or Poisson generalized linear model, with confounding factors such as space use, sightings frequency, or joint gregariousness as predictors. The residuals of this model are then used as measures of affiliation, as they reflect the difference between observed and expected association rates given the confounding factors. While a flexible and appealing approach, GAIs require that potential confounds be properly specified in terms of dyadic covariates, and that the relationship between confounds and edge weights be linear. This second issue, however, may be addressed by fitting generalized additive models (GAMs), where relationships are represented by smooth functions.

A related, but slightly different approach would be to incorporate confounds in the inferential model itself. Rather than deriving new edge weights via GAIs, if researchers identify likely confounds and summarize them quantitatively for each dyad or individual, these could be used directly in the final model. Where potential non-linearity between confounds and responses exist, data transformations, polynomials, and smooth functions may present a possible solution.

We feel that these methods have the potential to address the current issue that we have identified and we strongly encourage new work to explore and validate these approaches. It is important to note that the methods we propose are only useful if the question of interest is about the structure of social affinity, rather than the empirical pattern of encounters between individuals. If, instead, researchers are interested in the actual rates of contact (as is the case in disease research and studies of social learning), this approach may not be appropriate. Extensions of recent work using hidden state modelling may be more appropriate for disentangling true association patterns when detections are potentially biased or imperfect (Gimenez et al., 2019).

### Building better null models

The problems we have identified here arise because the commonly used null models for animal societies do not generate datasets representing the null hypothesis of interest in a regression setting. These models were specifically designed to test the null hypothesis of random social structure, not the null hypothesis that aspects of social structure are unrelated to exogenous factors. An obvious way forward would be the development of permutation procedures that generate datasets that correctly represent the relevant null hypothesis. In the case of dyadic regression, these datasets would maintain the structure of the data (e.g. sightings per individual, associations per sampling period, spatial patterns of observations), randomise identities of associated individuals, and simultaneously preserve the variance in edge weights. In the case of nodal regression, permuted datasets would maintain the same (or at least a similar) distribution of individual centrality within the network, in addition to structural confounds such as the size of groups, sightings per individual, and timing of sightings. The design of such procedures is far from trivial, and is beyond the scope of this paper, but we suspect that the development of algorithms that simultaneously maintain aspects of data structure and features of the social system will be an important area of methodological research going forward.

## Conclusion

The development of permutation techniques that control for sampling biases while maintaining temporal, spatial, and structural aspects of the raw data is an important development in the study of animal social systems, and we suspect that these procedures will remain a key tool for hypothesis testing in ecology and evolution. However, a lack of consideration regarding the matching up of the null hypothesis being tested with the null model being generated using datastream permutations has led to unwarranted application of these techniques, particularly in the context of hypothesis testing using regression models.

We recommend that researchers think critically and carefully about the null hypothesis they wish to test using social network data, and ensure that the null model they specify does in fact represent that hypothesis. We suspect that in most cases, the null hypothesis of random social structure will clearly not be appropriate, and therefore traditional datastream permutations will not be a viable approach. We hope that our discussion of this issue and the results of our simulations will result in reconsideration of how researchers employ null models when analysing animal social networks, promote further research and discussion in this area, and lead to the development of procedures that correctly specify null hypotheses and allow robust inference in animal social network studies.

## Supporting information

Simulation code

## Acknowledgements

We would like to thank colleagues at the Centre for Research in Animal Behaviour, particularly the members of the CRAB Social Network Club, for extremely valuable discussion that greatly improved this manuscript. DPC, DWF and SE acknowledge funding from NERC (NE/S010327/1). LJNB acknowledges funding from the NIH (R01AG060931; R01MH118203).

## Author contributions

MNW conceived of the project and designed the simulations, with input from all authors. All authors contributed to drafting the manuscript.

## Data availability

This study used no empirical data. All code necessary to reproduce our analysis is included in the online supplementary material.

## References

Anderson, M. J., & Robinson, J. (2001). Permutation tests for linear models. Australian and New Zealand Journal of Statistics, 43(1), 75–88. https://doi.org/10.1111/1467-842X.00156

Bejder, L., Fletcher, D., & Bräger, S. (1998). A method for testing association patterns of social animals. Animal Behaviour, 56(3), 719–725. https://doi.org/10.1006/anbe.1998.0802

Cowl, V. B., Jensen, K., Lea, J. M. D., Walker, S. L., & Shultz, S. (2020). Sulawesi Crested Macaque (Macaca nigra) Grooming Networks Are Robust to Perturbation While Individual Associations Are More Labile. International Journal of Primatology, 41(1), 105–128. https://doi.org/10.1007/s10764-020-00139-6

Croft, D. P., Madden, J. R., Franks, D. W., & James, R. (2011). Hypothesis testing in animal social networks. In Trends in Ecology and Evolution (Vol. 26, Issue 10, pp. 502–507). Elsevier Current Trends. https://doi.org/10.1016/j.tree.2011.05.012

Dekker, D., Krackhardt, D., & Snijders, T. A. B. (2007). Sensitivity of MRQAP tests to collinearity and autocorrelation conditions. Psychometrika, 72(4), 563–581. https://doi.org/10.1007/s11336-007-9016-1

Farine, D. R. (2017). A guide to null models for animal social network analysis. Methods in Ecology and Evolution, 8(10), 1309–1320. https://doi.org/10.1111/2041-210X.12772

Farine, D. R., & Whitehead, H. (2015). Constructing, conducting and interpreting animal social network analysis. Journal of Animal Ecology, 84(5), 1144–1163. https://doi.org/10.1111/1365-2656.12418

Firth, J. A., Sheldon, B. C., & Brent, L. J. N. (2017). Indirectly connected: simple social differences can explain the causes and apparent consequences of complex social network positions. Proceedings of the Royal Society B: Biological Sciences, 284(1867), 20171939. https://doi.org/10.1098/rspb.2017.1939

Franks, D. W., Ruxton, G. D., & James, R. (2010). Sampling animal association networks with the gambit of the group. Behavioral Ecology and Sociobiology, 64(3), 493–503. https://doi.org/10.1007/s00265-009-0865-8

Gimenez, O., Mansilla, L., Klaich, M. J., Coscarella, M. A., Pedraza, S. N., & Crespo, E. A. (2019). Inferring animal social networks with imperfect detection. Ecological Modelling, 401, 69–74. https://doi.org/10.1016/j.ecolmodel.2019.04.001

Hinde, R. A. (1976). Interactions, Relationships and Social Structure. Man, 11(1), 1–17.

James, R., Croft, D. P., & Krause, J. (2009). Potential banana skins in animal social network analysis. In Behavioral Ecology and Sociobiology (Vol. 63, Issue 7, pp. 989–997). Springer. https://doi.org/10.1007/s00265-009-0742-5

Krause, S., Mattner, L., James, R., Guttridge, T., Corcoran, M. J., Gruber, S. H., & Krause, J. (2009). Social network analysis and valid Markov chain Monte Carlo tests of null models. Behavioral Ecology and Sociobiology, 63(7), 1089–1096. https://doi.org/10.1007/s00265-009-0746-1

Kurvers, R. H. J. M., Krause, J., Croft, D. P., Wilson, A. D. M., & Wolf, M. (2014). The evolutionary and ecological consequences of animal social networks: Emerging issues. In Trends in Ecology and Evolution (Vol. 29, Issue 6, pp. 326–335). Elsevier Ltd. https://doi.org/10.1016/j.tree.2014.04.002

Manly, B. F. J. (1997). Randomization, bootstrap, and Monte Carlo methods in biology (2nd ed.). Chapman & Hall.

Miklós, I., & Podani, J. (2004). Randomization of presence-absence matrices: Comments and new algorithms. Ecology, 85(1), 86–92. https://doi.org/10.1890/03-0101

Poirier, M. A., & Festa-Bianchet, M. (2018). Social integration and acclimation of translocated bighorn sheep (Ovis canadensis). Biological Conservation, 218, 1–9. https://doi.org/10.1016/j.biocon.2017.11.031

Psorakis, I., Roberts, S. J., Rezek, I., & Sheldon, B. C. (2012). Inferring social network structure in ecological systems from spatiotemporal data streams. In Journal of the Royal Society Interface (Vol. 9, Issue 76, pp. 3055–3066). Royal Society. https://doi.org/10.1098/rsif.2012.0223

Psorakis, I., Voelkl, B., Garroway, C. J., Radersma, R., Aplin, L. M., Crates, R. A., Culina, A., Farine, D. R., Firth, J. A., Hinde, C. A., Kidd, L. R., Milligan, N. D., Roberts, S. J., Verhelst, B., & Sheldon, B. C. (2015). Inferring social structure from temporal data. Behavioral Ecology and Sociobiology, 69(5), 857–866. https://doi.org/10.1007/s00265-015-1906-0

Ryder, T. B., Horton, B. M., van den Tillaart, M., Morales, J. D. D., & Moore, I. T. (2012). Proximity data-loggers increase the quantity and quality of social network data. Biology Letters, 8(6), 917–920. https://doi.org/10.1098/rsbl.2012.0536

Spiegel, O., Leu, S. T., Sih, A., & Bull, C. M. (2016). Socially interacting or indifferent neighbours? Randomization of movement paths to tease apart social preference and spatial constraints. Methods in Ecology and Evolution, 7(8), 971–979. https://doi.org/10.1111/2041-210X.12553

Webber, Q. M. R., & vander Wal, E. (2019). Trends and perspectives on the use of animal social network analysis in behavioural ecology: a bibliometric approach. Animal Behaviour, 149, 77–87. https://doi.org/10.1016/j.anbehav.2019.01.010

Whitehead, H. (1999). Testing association patterns of social animals. Animal Behaviour, 57(6), F26–F29. https://doi.org/10.1006/anbe.1999.1099

Whitehead, H. (2008). Precision and power in the analysis of social structure using associations. Animal Behaviour, 75(3), 1093–1099. https://doi.org/10.1016/j.anbehav.2007.08.022

Whitehead, H., Bejder, L., & Ottensmeyer, C. A. (2005). Testing association patterns: Issues arising and extensions. Animal Behaviour, 69(5). https://doi.org/10.1016/j.anbehav.2004.11.004

Whitehead, H., & James, R. (2015). Generalized affiliation indices extract affiliations from social network data. Methods in Ecology and Evolution, 6(7), 836–844. https://doi.org/10.1111/2041-210X.12383

Zeus, V. M., Reusch, C., & Kerth, G. (2018). Long-term roosting data reveal a unimodular social network in large fission-fusion society of the colony-living Natterer’s bat (Myotis nattereri). Behavioral Ecology and Sociobiology, 72(6), 1–13. https://doi.org/10.1007/s00265-018-2516-4

